# Confidence and insight into working memories are shaped by attention and recent performance

**DOI:** 10.1101/2024.08.09.607293

**Authors:** Sammi R Chekroud, Anna C Nobre, Nils Kolling

## Abstract

Working memory (WM) is capacity-limited, and our ability to access information from WM is variable, but selective attention to working memory contents can improve performance. People are able to make introspective judgements regarding the quality of their memories, and these judgements are linked to objective memory performance. However, it remains unknown whether benefits of internally directed attention on memory performance occur alongside commensurate changes in introspective judgments. Across two experiments, we used retrospective cues (retro-cues) during working-memory maintenance to direct attention to items in memory. We then examined their consequence on introspective judgements. In the second experiment, we provided trial-wise feedback on performance. We found that selective attention improved confidence judgements and not just performance of the probed item. We were also able to judge participants’ genuine insight into working-memory contents through the correlation between confidence judgements and memory quality. Neurophysiologically, alpha desynchronization correlated first with memory error and then confidence during retro-cueing, suggesting a sequential process of attentional enhancement of memory contents and introspective insight. Further, we showed that participants can use feedback on the accuracy of confidence judgements to update their beliefs across time, according to performance. Our results emphasize flexibility in working memory by showing we can selectively modulate our confidence about its contents based on internally directed attention or objective feedback.

## Introduction

Working memory (WM) describes a core cognitive ability to store and manipulate information on short time scales to guide behaviour. It plays an important role in action planning, decision-making, and other complex behaviours. Its use is ubiquitous throughout daily life; from rummaging for a top to match the trousers you picked out to wear, to holding online the objects to collect before leaving the house in the morning.

However, not all memories should guide behaviour equally. WM is fundamentally limited (Bays & Husain, 2008; Fougnie et al., 2012; Luck & Vogel, 1997; Zhang & Luck, 2008) and the quality of our WM recall fluctuates across trials within tasks. As such, we cannot simply rely on memory at all times. Given this limitation, to be more adaptive and support optimal behaviour, the uncertainty of memories should also play an important part in their integration into various behaviours. When uncertainty regarding the contents of a memory is high, one should be wary of using it to guide behaviour (as we may make an incorrect, or suboptimal behaviour). When uncertainty regarding the contents of a memory is low, we can more appropriately integrate such memories into behaviour.

Despite the importance of knowing the reliability of internal representations for behaviour, uncertainty in WM is not typically studied. However, there is evidence that individuals have insight into the quality of their WM representations. Using categorical confidence judgements, it has been shown that both prospective and retrospective confidence judgements vary systematically with response errors (Rademaker et al., 2012). Low confidence in WM was associated with increased errors and higher guessing rates. Systematic manipulation of WM load (three vs six items to maintain in memory) also revealed lower reported confidence (both prospectively and retrospectively) when load was high. Even when only one item is held in WM, the same association between high confidence and high accuracy (low error) has been found. This confidence in WM also appears to influence interactions between trials, where items reported with higher confidence exert stronger biases on subsequent trials than items reported with lower confidence (Samaha et al., 2019).

Moving beyond categorical judgements, others have investigated whether and how uncertainty may be parametrically related to WM accuracy (Honig et al., 2020; Li et al., 2021; Yoo et al., 2021). WM tasks often use precision-recall tasks in which participants are required to report the exact value of a feature of an object held in working memory (Bays et al., 2009; Zhang & Luck, 2008). Rather than producing categorical confidence estimates, such tasks have been adapted to allow individuals to report their precise uncertainty in WM contents in the same circular space (Honig et al., 2020; Li et al., 2021). This has revealed a parametric relationship between accuracy and subjective confidence. Permutation-based analyses have shown that participants do not simply use heuristics about colour difficulty to guide such confidence judgements, as the observed relationship between accuracy and confidence was stronger than simulated relationships based on feature similarity (Honig et al., 2020). As a result, confidence judgements concerning memorised items appear not to be based solely on external elements such as heuristics or stimulus features, but instead appear to be based (at least in part) on internal mnemonic representations themselves.

However, not all contents in working memory are represented equally. Attention can be directed to specific information held in WM to improve task performance (Griffin & Nobre, 2003; Landman et al., 2003; Stokes & Nobre, 2012). Retrospective attention-directing cues (‘*retrocues*’) to locations (Griffin & Nobre, 2003; Landman et al., 2003; Myers et al., 2018), objects, and features (Hajonides et al., 2020; Niklaus et al., 2017) of items maintained in WM can improve performance on tasks. To measure the quality of WM contents and associated benefits of attention-directing cues, WM studies typically report variable precision of responses across trials, and this variability serves as a proxy readout of fluctuations in the precision of encoding or maintenance of the probed items. Such attentional manipulations typically show that the width of response error distributions for the recalled features are more precise (reduced variance) when items are held in a prioritised attentional state.

Knowing that WM precision varies, not all working memories should guide behaviour equally. Previous studies have shown that confidence judgements appear to track endogenous variability in WM performance (Honig et al., 2020), suggesting these judgements tap into the quality of internal representations. However, it is currently unknown whether such introspective confidence judgements can track functional changes in the attentional state of WM contents. If the subjective uncertainty relating to an object held in WM changes based on its attentional state, it would more convincingly demonstrate that such introspective judgements are tracking the state of the internal mnemonic representation itself.

The current study aimed to address two important questions. Firstly, does internal attention change introspection about WM quality in tandem with objective improvements in performance? If introspective judgements tap into the quality of internal representations, attentional manipulations that have been shown to alter the state of internal representations should also affect such introspective judgements. And if it does, what are the neural processes changing with increased confidence based on attentional shifts? Secondly, does feedback concerning introspective ability lead to adaptive changes in these introspective responses? It is known that there are biases between trials based on stimulus history, and such biases are influenced by the confidence with which an item was encoded (Samaha et al., 2019). Are introspective judgements similarly influenced by the accuracy of our previous introspections?

To address these questions, we performed two experiments. In the first experiment, participants reproduced the orientation of a bar previously encoded into working memory. On half of all trials, a retrocue was presented during the maintenance delay indicating which item was to be reported on that trial. Following orientation reproduction in each trial, we introduced a novel confidence-judgement probe for participants to report the subjective uncertainty in their response. In a second experiment, we concurrently recorded electroencephalographic (EEG) measurements. Given previous studies have shown that introspective confidence judgements are systematically related to WM precision, we aimed to test whether neural markers of attentional selection within WM were parametrically associated with the introspective judgement and whether attentional effects on accuracy are dissociable from confidence changes. Further, we included quantitative feedback on every trial. This feedback showed individuals their response error and their degree of confidence, allowing an evaluation of the accuracy of their confidence judgement. This allowed us to explore behaviourally whether and how feedback can act as a learning signal to shape confidence judgements across trials.

In both experiments, we used a precision-recall working-memory task in which participants were probed to report the orientation of one of two items from a to-be-remembered array. After report, participants made a confidence judgement on the same circular space – reporting a range of angles centred upon their initial response to indicate how confident they were in their reproduction. In line with previous reports, we observed that retrocue-based attentional prioritisation improved working-memory precision. Importantly, we observed that retrocues also improved introspective judgements. We show that attentional cues improved the accuracy of the introspective judgement made on each trial, showing that attentional cues can also change the quality of insight into our internal representations. Using EEG, we show that the degree of retrocue-induced alpha lateralisation, across trials, is first predictive of the accuracy of recall from WM, and subsequently of the degree of confidence in WM recall, at two temporally-distinct stages. Importantly, This suggests that confidence is sensitive to changes in the attentional state of the internal representation, but attentional benefits on the objective and subjective aspects of the WM representation are at least partially distinct neural processes. Further, we demonstrated that participants can use feedback about the error in their confidence judgements (how over- or under-confident they were) to flexibly adjust their confidence across trials. Overall, this shows that while participants have some access to their internal representations (and that access can be boosted with internal attention), external factors, such as feedback, are also flexibly incorporated to gain a more accurate insight into the quality of working memories.

## Methods – Experiment 1

Experimental procedures for both experiments were reviewed and approved by the Central University Ethics Committee of the University of Oxford. All participants provided written, informed consent before they participated in the study.

### Participants

Twenty-two participants were recruited for Experiment 1 (7 males, 15 females). Two were excluded from data analysis (one failed to confirm their response in almost 40% of trials, a second responded randomly across trials), leaving twenty data sets that were fully analysed (6 male, 14 female, age range 20-35 years, mean age 26.9 years, 1 left-handed). All participants reported having normal or corrected-to-normal vision and not being colour blind. Participants were compensated £10 per hour for their time. Participants completed 8 blocks of 32 trials in one session, resulting in a total of 256 trials.

### Stimuli, procedure, and task

Participants were seated in a dimly lit room approximately 40 cm from the monitor (24-inch; resolution: 1920 x 1080 pixels; screen width: 53 cm; refresh rate: 100 Hz). The experimental script was generated using PsychoPy version 1.90.3 (Peirce et al., 2019).

Figure 1 shows a schematic of the task. At the start of each trial, two coloured bars appeared either side of a central fixation cross (RGB: [150, 150, 150]), for 250 ms. The centre of each bar was six degrees of visual angle (dva) from the centre of the screen, with a height and width of 5.7 and 0.8 dva respectively. Bars could either be blue (RGB: [0, 0, 255]) or yellow (RGB: [255, 255, 21]). Orientations were drawn independently from uniform distributions between 0 and 179 degrees.

**Figure 1.**
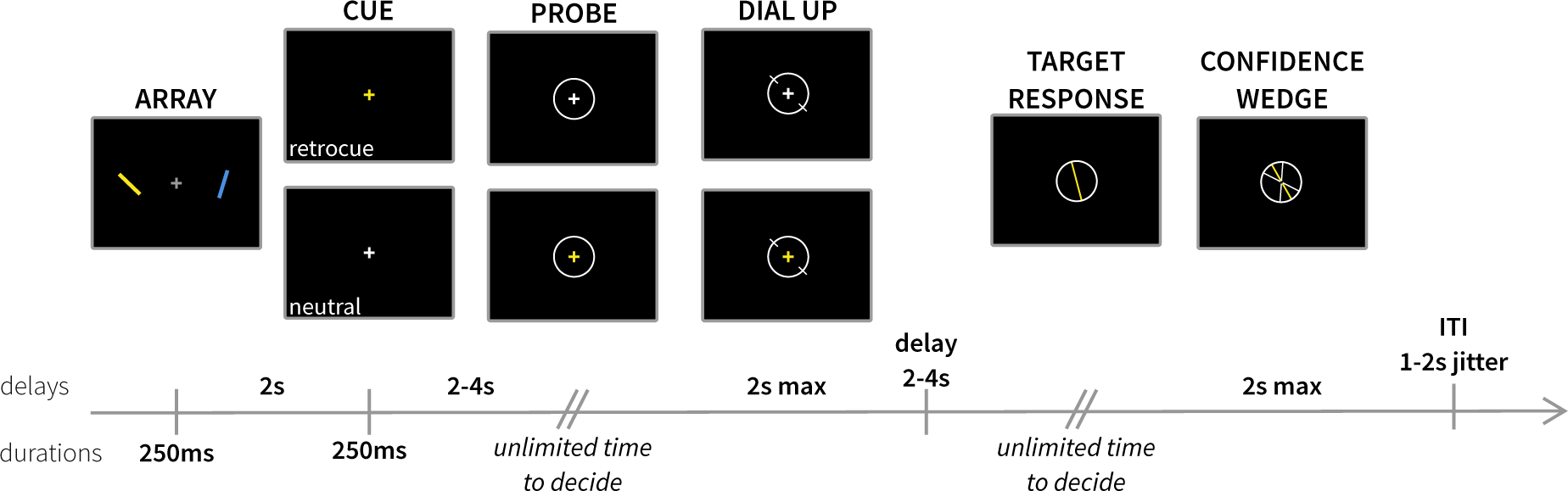
A) Trial structure for experiment 1. Participants were presented with two coloured, oriented bars to memorise. After a delay, a cue was presented that was either informative to which item they would be probed about (retro-cue) or uninformative (neutral cue). Participants subsequently reproduced an orientation from memory. After reporting an orientation, participants reported their confidence in their response. This was done by creating a wedge that reflected a range of values they believed the target orientation could be within – larger wedges reflect lower confidence in their response.

After a two-second delay, the fixation cross changed colour for 250 ms. On 50% of all trials, the fixation cross changed colour to match one of the items previously encoded into memory, signalling that only this item was relevant for subsequent report (*retrocue* condition). In the other half of trials, the fixation cross changed to white, providing no information regarding item prioritisation (*neutral* condition). After the cue, there was a variable delay between 2 and 4 seconds (drawn randomly from a uniform distribution in 100-ms steps) before the probe stimulus appeared on screen. In neutral trials, the fixation cross changed colour to match the item that participants had to report. On retrocue trials, the fixation cross changed to white, and participants reported the previously cued item (thus ensuring participants processed the cue). A circle appeared around the fixation cross (diameter: 5.7 dva) to indicate the start of the probe phase. Participants had unlimited time to decide on the orientation to report before pressing the ‘space’ bar (with their left hand) to indicate they were ready. Upon pressing the space bar, two white lines appeared on opposite sides of the probe circle. The starting orientation for the dial was randomly drawn from a uniform distribution between 0 and 179 degrees. Participants used the mouse to rotate the dial around the circle to replicate the orientation of the probed item. A left mouse click was required to confirm their response. Although participants had an unlimited time to decide on the orientation to report, they were limited to two seconds to deliver and confirm their orientation response. This reporting method therefore provides informative measures relating to accessing and deciding on the item to report as well as to the contents of items reported (see also Niklaus et al., 2017; van Ede et al., 2018, 2019).

After clicking to confirm the response, there was a variable delay between 2 and 4 s before the confidence-report phase began. Participants were presented with their orientation response as a bar inside a circle, with the colour matching the probed item. Participants had unlimited time to decide how confident they were in the response they gave, before pressing the ‘space’ bar with their left hand to initiate their response. Upon pressing the space bar, two white lines appeared, mirrored symmetrically around the response orientation. Using the mouse, participants adjusted the angles formed by these lines to form a wedge to indicate their confidence interval in a circular space. Participants were instructed to use the circular space, with larger wedges corresponding to low confidence in their response and narrower wedges indicating high levels of confidence in the orientation they reproduced. Participants had up to two seconds to dial and confirm their confidence report before the trial ended. After responding, there was an inter-trial interval between 1 and 2 s (randomly drawn from a uniform distribution in 100-ms steps) before the next trial began, during which the grey fixation cross was presented.

### Behavioural Analysis

We defined two main sources of error. *Response error* (see Figure 2A) was calculated as the angular deviation between the target orientation and the response. *Confidence error* (see Figure 3A) was calculated as the difference between this response error and the reported confidence wedge on any given trial (i.e. the discrepancy between error and confidence). The time between probe onset and pressing ‘space’ to start the dial phase was taken as the decision time. Where appropriate to analyse aggregate data, repeated-measures Analysis of Variances (ANOVAs) were used. Repeated-measures ANOVAs and random-effects analyses implementing generalised linear models (GLMs) were conducted using the *statsmodels* package (version 0.14.0) in python (Seabold & Perktold, 2010).

**Figure 2.**
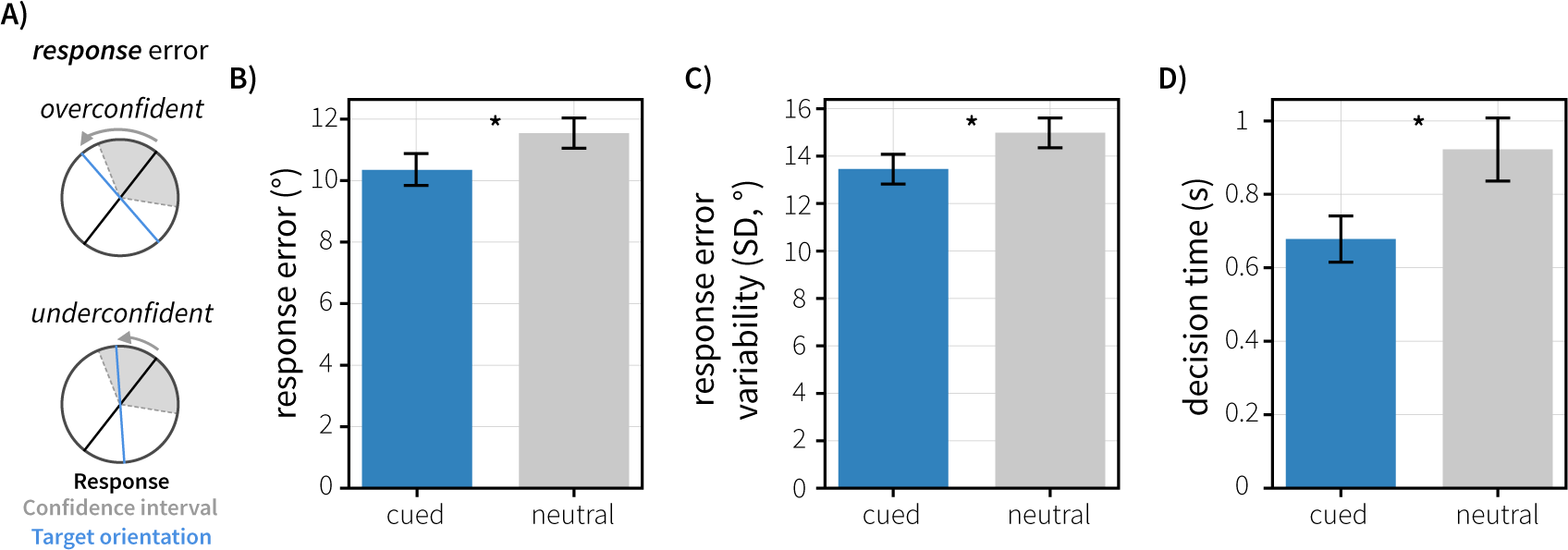
Behavioural performance in Experiment 1. A) Schematic visualising response error (deviation between response and target) in this task. Confidence wedges (grey shaded area) are centred on the response given by the participant (black line). The blue line represents the originally presented target orientation. B) Mean absolute response error was modulated by Attention. C) Average response error variability (standard deviation of response errors in degrees) for cued and neutral trials. D) Decision time (time taken to initiate the orientation reproduction) was modulated by cueing. All errorbars reflect the standard error of the mean.

**Figure 3.**
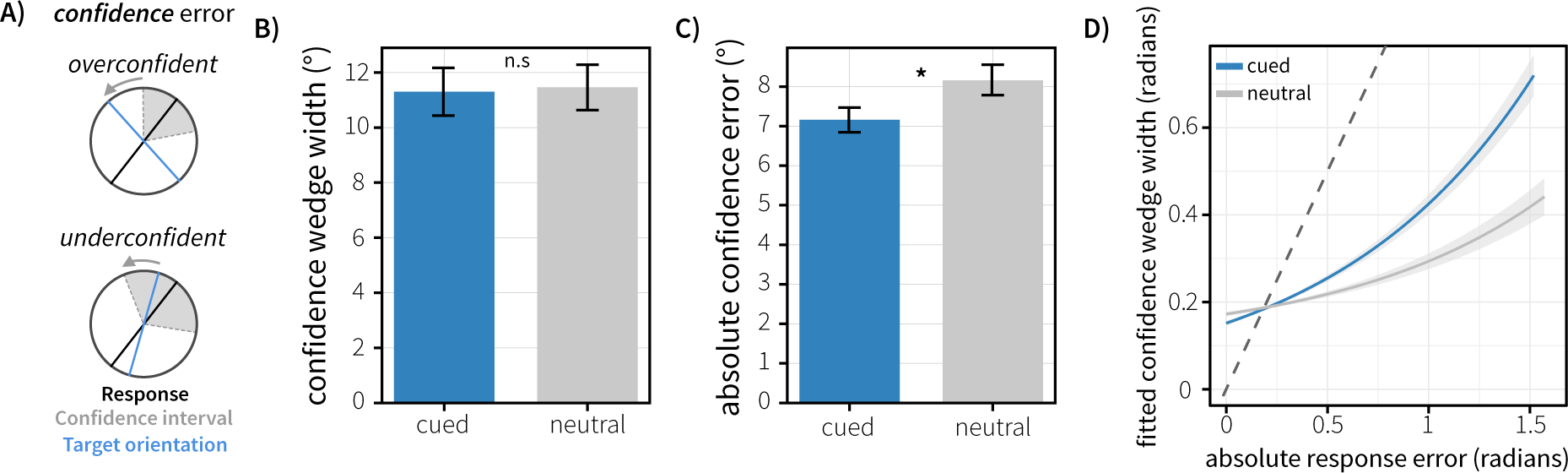
Behavioural performance in Experiment 1. A) Schematic illustration of confidence error for trials where participants were overconfident or underconfident. B) Average confidence wedge width (degrees) for cued and neutral trials.. C) Average confidence error for cued and neutral trials. D). Confidence wedge width is parametrically associated with response error across trials. Confidence wedge width was modelled as a function of response error, cue type (Neutral vs Cued) and their interaction, using linear-mixed effects modelling. Fitted confidence is visualised against response error, controlling for across-subject variance in effects. The dashed line represents the identity line, which would indicate perfect insight into memory quality. All error bars and shaded error-regions represent the standard error of the mean.

Two types of analyses were used to test the relationship between response error and subjective confidence in working memory and the influence of attention on this relationship. First, we implemented a random-effects analysis. We fit a GLM to the behavioural data separately for each subject. This model predicted the logarithm of the reported confidence wedge width (to approximate a normal distribution) as a function of condition (cued vs neutral), response error, and the interaction term for these two. Attention condition was estimated as a contrast regressor to compare the difference between cued vs. neutral trials. An intercept term was included to capture individual subject variance, and all regressors were standardised (z-scored) prior to model fitting. We then subjected the single-subject beta coefficients to an across-subject t-test to test for significance of the effect at the group level.

Secondly, we fit a linear mixed-effects model (LMM) using the *‘lmerTest’* package in R (Kuznetsova et al., 2017) to provide p-value estimates for coefficients. LMMs take into account all single-trial variance rather than aggregating data and allow us to look at the strength and direction of relationships while statistically controlling for other correlated effects. To test which factors are linked to confidence judgements using this LMM approach, we modelled confidence as a function of response error, Attention (neutral vs cued), and the interaction term for these two. We initially chose a maximal random-effects structure that included random intercepts for error, condition, their interaction, and participant. However, maximal random-effects structures can lead to over-parameterisation of models and high degrees of freedom during model fitting. To have our models converge on more stable solutions, we used a Principal Component Analysis (PCA) of the random-effects variance-covariance estimate for the fitted mixed-effects models to identify over-parameterisation (as in Lauer et al., 2018). Random slopes that were not supported by the model and did not contribute significantly to the goodness of fit were removed.

## Results – Experiment 1

Behavioural results are visualised in Figures 2 and 3. To assess whether retrocues affected behaviour in our task, we conducted one-way repeated-measures ANOVAs on decision time, response error, response variability, and confidence. We found a main effect of Attention on decision time (*F*(1, 19) = 21.41, *p* < 0.001, *ƞ*^2^_*p*_ = 0.530) and response error (*F*(1, 19) = 10.66, *p* = 0.004, *ƞ*^2^_*p*_ = 0.359). Responses were faster in cued trials (*M* = 0.678s, *SE* = 0.063s) than neutral trials (*M* = 0.922s, *SE* = 0.0857s) and were more accurate in cued trials (*M* = 10.3 degrees, *SE* = 0.504 degrees) than neutral trials (*M* = 11.5 degrees, *SE* = 0.493 degrees). We also found a main effect of Attention on response variability (*F*(1,19) = 10.1, *p* = 0.00499, *ƞ*^2^_*p*_ = 0.347), with lower variability (lower circular standard deviation) in the response distribution for cued trials (*M* = 13.4 degrees, *SE* = 0.625 degrees) than neutral trials (*M* = 15.0 degrees, *SE* = 0.625 degrees).

A key facet of this task was the inclusion of a confidence judgement on each trial of the experiment. Participants were asked to report their confidence in the working-memory response they had just made (their initial orientation reproduction). Critically, this judgement was made in the same circular space, and participants were instructed that a narrow wedge would indicate high confidence, with a large wedge indicating low confidence in their response. To test whether response error and confidence were related across trials (and ensure participants were using the confidence judgement appropriately), we implemented two complementary analyses: a random-effects and a linear-mixed effects model analysis, to assess the relationship between the response error and confidence.

The random-effects analysis revealed a significant relationship between response error and confidence. A t-test on the single-subject beta coefficients showed a significant effect of response error (*t*(19) = 6.41, *p* = 3.81*10^-6^). Trials with higher response errors were associated with larger reported confidence wedges. There was also a significant interaction between response error and condition (*t*(19) = 3.09, *p* = 0.00599). This interaction demonstrates that the relationship between response error and confidence was stronger (more positive beta) for cued trials compared to neutral trials, across subjects.

The LMM analysis highlighted an association between response error and confidence wedge widths across trials (see Figure 3D). Trials with higher response errors were associated with larger confidence wedge widths (*β* = 0.651, *t* = 6.913, *p* = 1.04*10^-6^). We also found that Attention (neutral or cued) interacted with this relationship (*β* = -0.136, *t* = -3.127, *p* = 0.00178), suggesting that the relationship between response error and confidence is influenced by selective attention during the maintenance period. This was driven by a stronger association between response error and confidence wedge size (more positive beta) in cued trials compared to neutral trials. To explore this further, we conducted one-way repeated-measures ANOVAs on confidence wedge width and confidence error. We saw no significant effect of cue type on confidence wedge widths (*p* = 0.459). However, there was a strong main effect of Cue on confidence error (*F*(1,19) = 25.54, *p* < 0.001, *ƞ*^2^_*p*_ = 0.573), with confidence error lower in cued trials (*M* = 7.16 degrees, *SE* = 0.313 degrees) than neutral trials (*M* = 8.16 degrees, *SE* = 0.396 degrees).

Our analyses are predicated on the idea that individuals have insight into their single-trial performance or memory quality. This single-trial insight is integrated into their estimate of subjective confidence, leading to a relationship between response error and confidence. An alternative hypothesis of WM confidence could argue that perfect insight into single-trial memory quality would give rise to precisely zero response error. For example, knowing that your memory representation is exactly six degrees inaccurate, an optimal agent would adjust their orientation reproduction by six degrees to accommodate this. Instead, one might have perfect insight into the across-trial response error distribution, and sample from this distribution to generate confidence estimates. To test whether our observed effects could arise under this hypothesis, we implemented control analyses. We first use a permutation-based analysis to show that empirically observed correlations between response error and confidence are unlikely to arise without insight into single-trial performance (see Supplementary Analysis 1, and Supplementary Figure 1). We then show, through both permutation of our observed data, and simulation of artificial data, that the findings of both our random-effects analysis (Supplementary Analysis 2, see Supplementary Figure 2 for visualisation) and our LMM analysis (Supplementary Analysis 3, see Supplementary Figure 3 for visualisation) are unlikely to arise under this alternative hypothesis. Instead, converging evidence from our observed empirical findings, simulations, and permutations of empirical data suggests that individuals have insight into WM representations. Further, this insight is influenced by the attentional state of the maintained representations.

## Methods – Experiment 2

### Participants

Twenty-six healthy human volunteers participated in the study (age range 18-36, mean age 25.2 years, 17 female, 2 left-handed). The study involved two task sessions separated by a break, each comprising of 8 task blocks of 32 trials, resulting in a total of 512 trials. Six participants were excluded prior to data analysis: five completed only one session of task, and an additional participant was removed due to excessive noise in the raw EEG data. This resulted in a final sample of 20 participants (age range 18-36 years, mean age 25.3 years, 14 female, 1 left-handed). All participants reported normal or corrected-to-normal vision and not being colour blind. All participants provided written informed consent prior to participation and were reimbursed £15 per hr for their time.

### Stimuli, procedure and task

Participants were seated in an unlit, electrically shielded booth 100 cm from the monitor (27-inch, resolution: 1920 x 1080 pixels, screen width: 60 cm, refresh rate: 100 Hz). The experimental script was programmed using PsychoPy version 1.90.3 (Peirce et al., 2019).

The task structure for Experiment 2 was similar to that of Experiment 1. However, in addition to the confidence judgement participants made, feedback relating to performance was presented on each trial. All stimulus durations remained the same, but some delays between events changed. Figure 4 shows the task schematic with stimulus timings. The delay between the array offset and cue onset was reduced to 1s. The delay between cue and probe was fixed at 1.5 s. The delay between response confirmation and the confidence report phase beginning was 1 – 1.5 s.

**Figure 4.**
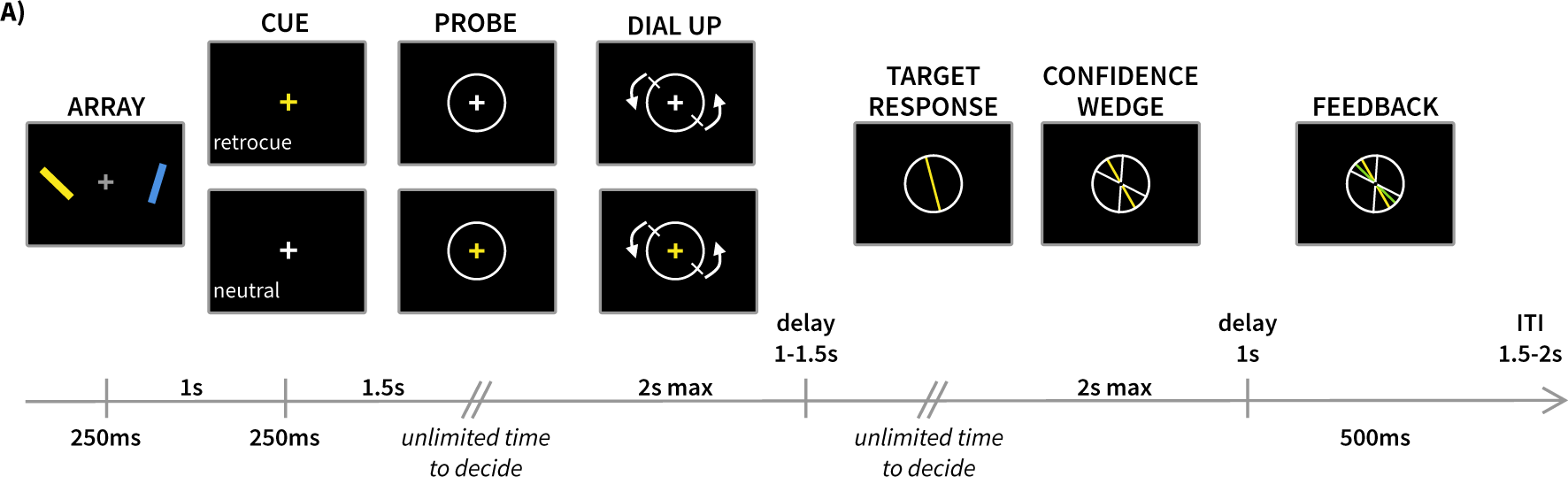
Trial structure in Experiment 2. A) Participants were presented with two coloured, oriented bars to memorise. After a delay, a cue was presented that was either informative (retro-cue) or uninformative (neutral cue) with respect to which item they would be probed about. Participants subsequently reported an orientation from memory. After reporting this orientation, they reported their confidence in their response, as in Experiment 1. After confirming their confidence response, participants were presented with feedback on their performance. Participants were presented with their responded orientation and their confidence wedge. The original target was then overlaid on top, coloured red or green depending on whether the target was outside or inside the reported confidence wedge (respectively). This gave binary information about the correctness of their confidence judgement, and continuous information regarding the accuracy of their orientation reproduction.

Uniquely to Experiment 2, a feedback routine was included on every trial after participants reported their confidence, providing comprehensive feedback on their response and confidence performance. After clicking to confirm their confidence judgement, the visual display remained fixed. If a participant clicked to confirm their confidence before the end of the two-second limit, their response remained on screen until the end of this limit, and a further one second before the feedback was presented. If a participant failed to confirm their response (i.e., it timed out after two seconds), there would still be a 1-s delay before feedback was presented. Participants were always shown their response and confidence width for at least one second. After this delay, the feedback was presented. Participants were shown the original target orientation as a thin bar (length: 5.8 dva, width: 0.1 dva, the same size and shape as the confidence interval boundary lines). This original orientation was coloured depending on the accuracy of the confidence judgement: if the target was outside the reported confidence interval, the bar was red (RGB: [204, 0, 0]); if the target was inside the interval, the bar was green (RGB: [0, 255, 0]). Therefore, the colour of the bar provided binary categorical correct/incorrect feedback about the confidence judgment, alongside the complete, continuous information available from the distance between the confidence width and the orientation of the feedback bar. This gave participants information with which to evaluate their own performance and determine whether the beliefs they held about their memory on a given trial were accurate (i.e. whether their confidence was justified). This allowed us to investigate whether participants could use this feedback information to adjust their confidence across trials. Feedback was presented for 500 ms before the visual display was replaced by the grey fixation cross for the duration of the ITI (1.5 – 2 s).

When a participant failed to confirm the initial orientation response within the two-second time limit, the last orientation on the screen was taken as the response and carried forward to the confidence phase to complete a trial. These trials were subsequently excluded from all analyses to ensure that only trials in which the participant explicitly reported an orientation were considered. At the end of each block, participants were provided with summary text informing them of their average reaction time (time taken to press the space bar to start the dial-up), their average response error (absolute angular distance between the target and response), their average tendency to be over- or under-confident, and the average degrees by which they were over- or under-confident.

### EEG acquisition and processing

EEG data were collected using a SynAmps RT amplifier and Curry 8 Software (Compumedics Neuroscan, North Carolina, USA) using a 61 Ag/AgCl sintered electrode array (EasyCap, Herrsching, Germany) according to the 10-10 montage layout. Data were referenced to the left mastoid during recording and re-referenced to an average-mastoid reference offline. The ground was placed on the upper left arm, and two bipolar electro-oculograms (EOG) were recorded. One pair was placed above and below the left eye to measure the vertical EOG and identify blinks, while the other was placed lateral of each eye to record the horizontal EOG and identify saccades. Data were sampled at 1000 Hz during acquisition and stored for subsequent analysis.

During preprocessing, data were converted from the native CDT format to SET format using the EEGLAB toolbox and custom code written in Matlab (version 2015b, Mathworks). EEG data were subsequently imported, pre-processed, and analysed in Python using the MNE-Python toolbox version 0.21 (Gramfort et al., 2013) and custom code. Raw data for each session were first band-pass filtered between 1 and 40 Hz. Denoising was then performed using Independent Component Analysis (ICA) using the ‘InfoMax’ routine for all EEG sensors. Components related to eye movements were detected by correlation with the horizontal and vertical EOG and subsequently removed from the data. Trials were discarded if blinks were detected around the array onset and cue onset period (as blinks in these periods would prevent processing of task-relevant information, and thus influence behaviour). An automated routine was implemented to detect blinks around these two events of interest. 500-ms epochs were taken from the EOG traces around these events, bandpass filtered between 1 and 40 Hz, and baselined to the 250 ms prior to the event. For each epoch, the difference between the peak and minimal voltage in the vertical EOG was calculated and z-scored across epochs. Epochs containing blinks were identified as those with z-scored difference values larger than two and the raw difference values greater than 250 microvolts. This was done separately for both cue and array events. Trial identifiers containing blinks were saved to mark trials to be excluded from further analyses. Data were subsequently epoched around events of interest, and trials were removed based on within-trial variance using an automated procedure that looks at the variance in the broadband signal at a significance threshold of 0.05 (generalised extreme studentized deviate test, GESD). This procedure was run separately for each event of interest. Trials were subsequently visually inspected to identify any remaining trials with excessive noise, for example in the baseline period, to be removed from analysis. Trials were only retained where participants explicitly confirmed their orientation reproduction response (clicking to confirm at the dial up phase) and decision time was within 2.5 standard deviations of the mean.

### Spectral analysis

Spectral power was estimated at frequencies using a multitaper decomposition (*tfr_multitaper* in MNE-python). For the cue-locked period, we took epochs from -0.5 to +2 s, to capture the entire maintenance period after cue presentation, and downsampled the data to 100 Hz. The single-trial power was estimated for frequencies between 1 and 40 Hz. For each frequency, we used a fixed 300-ms time window such that the number of cycles changed with the frequency being estimated. Where necessary to do so, time-frequency resolved data were baselined between -2 and -1.5 s prior to the event of interest (functionally equivalent to a 500-ms period prior to the presentation of the array, at the start of the trial).

For analysis of posterior alpha lateralisation, data were averaged over a cluster of electrodes in each hemisphere (PO7, PO3, O1; PO8, PO4, O2). Data were averaged across these two electrode clusters separately, then averaged across the alpha frequency range (8-12 Hz). This gives rise to a time course of alpha power in these two lateralised clusters of electrodes. Lateralisation was calculated relative to the side of the probed item on the trial and expressed as a normalised percentage [(contra-ipsi)/(contra+ipsi) * 100].

### Statistical analysis

GLMs were used to investigate stimulus-evoked alpha lateralisation and its relationship with behaviour. For statistical inferences, the output beta-weights for contrasts of interest were used as the input for cluster-based permutation testing. To aid interpretation of this analysis, we took the negative of absolute response error, such that it represented accuracy (with larger values reflecting increased accuracy on a trial). Confidence wedge width was also inverted such that it represented confidence (with larger values reflecting higher confidence in WM recall).

We tested for significance using cluster-based permutation statistics (all permutations, alpha = .05), which avoids the multiple-comparisons problem. On time-series data, this generates a null distribution by randomly permuting the trial-average data at the participant level along the time dimension. This was implemented using MNE-python’s 1 sample permutation cluster test to test the observed group-level timeseries against zero. Clustering was performed on the first-level beta weights for the contrast of interest, allowing us to investigate behavioural correlations with the data while controlling for correlated variables.

### Latency Analysis

We evaluated latency differences in beta coefficient timecourses by implementing a Jackknife analysis (Miller et al., 1998). This approach estimates a temporal standard error for the difference in onset latency for peaks in two timeseries. To do so, we iteratively removed one participant from the participant pool and estimated the point at which the beta coefficient timecourse reached 70% of its peak value. In each iteration, we calculated this onset point for the accuracy and confidence coefficient timecourses separately, and quantified the difference in onset time. Across iterations, this builds a distribution of the temporal difference in the onset of the peaks in the two timeseries, alongside a jackknife-based estimate of the standard error. We then formally test this distribution against zero to determine the statistical significance of any onset latency difference.

## Results – Experiment 2

To assess whether retrocues affected behaviour, we conducted one-way repeated-measures ANOVAs on decision time, response error, response variability, and confidence wedge width. The behavioural results are depicted in Figures 5, 6 and 7. We found a main effect of Attention on decision time (*F*(1,19) = 66.3, *p* < 0.001, *ƞ*^2^_*p*_ = 0.777) and response error (*F*(1,19) = 16.6, *p* < 0.001, *ƞ*^2^_*p*_ = 0.467). Responses were faster in cued trials (*M* = 0.529 s, *SE* = 0.0415 s) than neutral trials (*M* = 0.775 s, *SE* = 0.0597 s) and were more accurate in cued trials (error: *M* = 9.22 degrees, *SE* = 0.539 degrees) than neutral trials (*M* = 10.4 degrees, *SE* = 0.700 degrees). We also found a main effect of Attention on response variability (*F*(1,19) = 13.0, *p* = 0.00186, *ƞ*^2^_*p*_ = 0.407), with less variability (lower circular standard deviation) in the response distribution for cued trials (*M* = 12.2, *SE* = 0.694) than neutral trials (*M* = 13.7, *SE* = 0.891).

**Figure 5.**
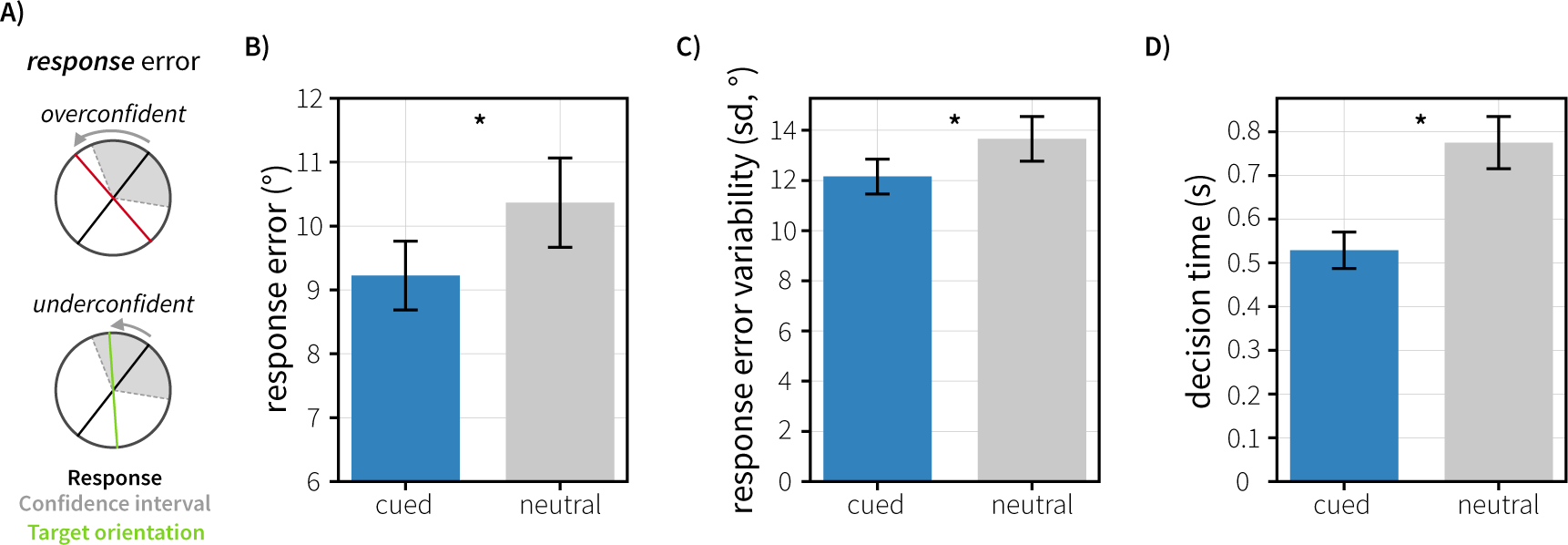
Behavioural performance in Experiment 2. A) Schematic visualising response error (deviation between response and target) in experiment 2. Confidence wedges (grey shaded area) are centred on the response given by the participant (black line). The red and green lines represent the originally presented target orientation, coloured by whether the confidence response was incorrect (original orientation outside of reported confidence wedge) or correct (original orientation inside the reported confidence wedge width). B) Mean absolute response error was modulated by Attention. C) Average response error variability (standard deviation of response errors in degrees) for cued and neutral trials. D) Decision time (time taken to initiate the orientation reproduction) was modulated by Attention. All errorbars reflect the standard error of the mean.

**Figure 6.**
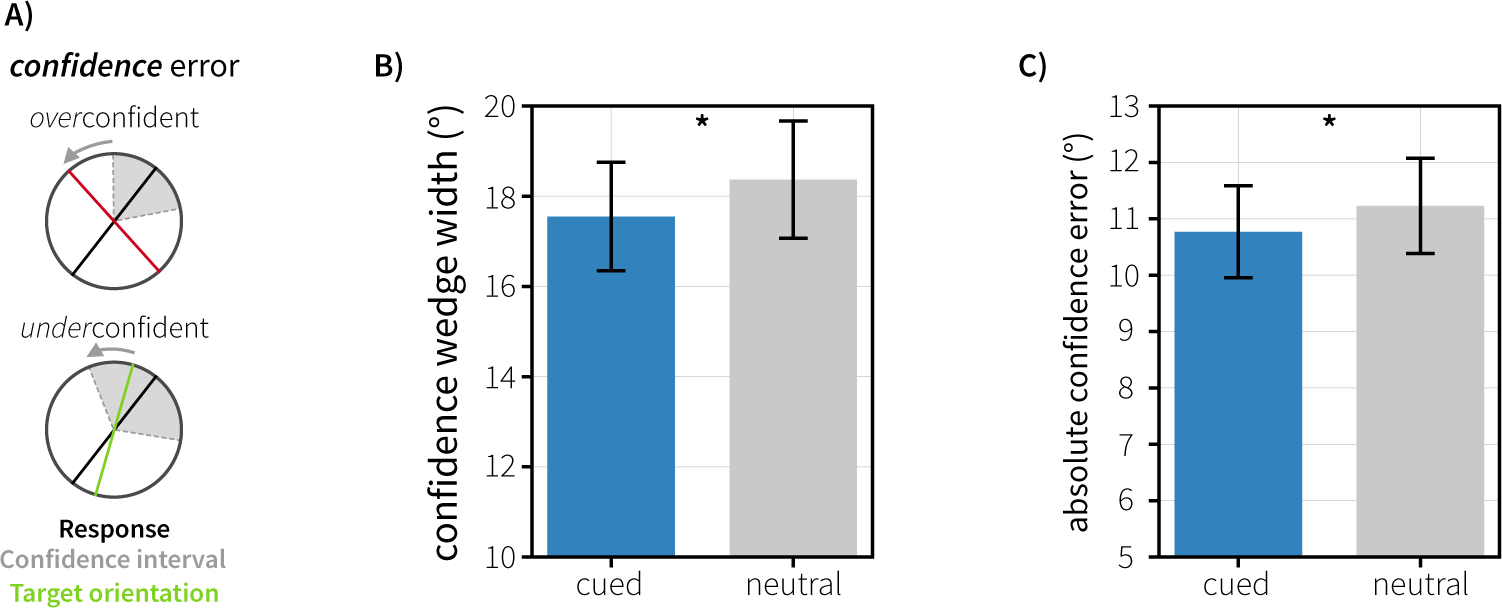
Behavioural performance in experiment 2. A) Schematic visualising confidence error and how it related to coloration of feedback. When the original target orientation was outside the reported confidence wedge, feedback was coloured red. When the target orientation was inside the reported confidence wedge, feedback was coloured green. B) Average confidence wedge width (degrees) for cued and neutral trials. C) Average absolute confidence error (how far away the target orientation was from the edge of the reported confidence wedge) for cued and neutral trials. Error bars reflect the standard error of the mean.

**Figure 7.**
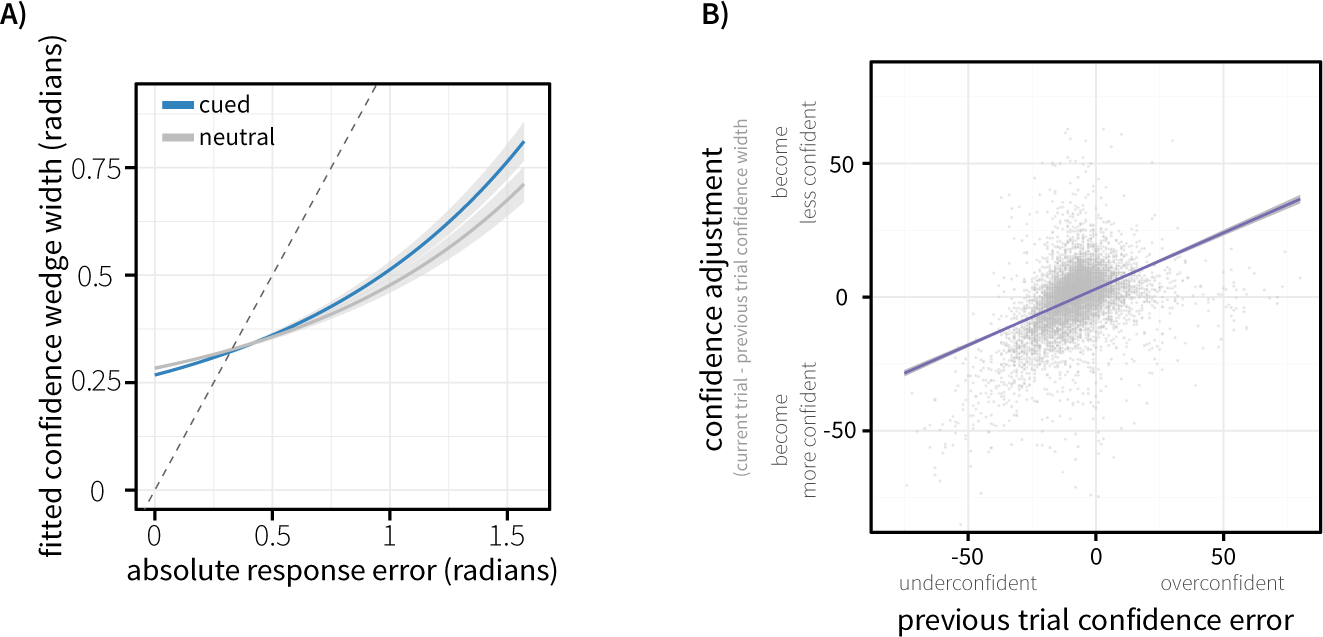
A) Relationship between absolute response error (radians) and confidence wedge width (radians), for cued and neutral trials. Confidence wedge width was modelled as a function of response error, cue type (neutral vs cued), and their interaction using linear mixed-effects modelling. Fitted confidence is visualised against response error, controlling for between-subject variance in effects. B) Visualisation of confidence updating across trials. Confidence error on the previous trial (degrees) is plotted against change in confidence between previous and current trial in degrees (i.e. whether confidence wedge widths increased or decreased based on confidence error on the previous trial).

When analysing the confidence data, we found a main effect of cue type on the reported confidence wedge width (*F*(1,19) = 17.8, *p* < 0.000459, *ƞ*^2^_*p*_ = 0.484). Confidence was higher in cued trials (i.e. narrower wedges; *M* = 17.5 degrees, *SE* = 1.20 degrees) than in neutral trials (*M* = 18.4 degrees, *SE* = 1.30 degrees).

As the confidence judgement was made in the same circular space as the orientation response, it was possible to test for the relationship between these two judgements across all trials, allowing us to look at the error of participants’ confidence judgements. As defined earlier, confidence error relates to the deviation between the original target orientation, and the edge of the confidence wedge reported on a trial. Due to how this value is computed, its sign holds meaning: positive values reflect over-confidence (target orientation was outside the confidence wedge) and negative values reflect under-confidence (target orientation was within the confidence wedge). The absolute confidence error therefore reflects the error in their confidence judgement, or how close they were to having perfect insight into their working memory. We found a main effect of Attention when looking at absolute confidence error (*F*(1,19) = 6.69, *p* = 0.0181, *ƞ*^2^_*p*_ = 0.260), with absolute confidence error lower in cued trials (*M* = 10.8 degrees, *SE* = 0.815 degrees) than neutral trials (*M* = 11.2 degrees, *SE* = 0.846 degrees).

A random-effects analysis showed a significant relationship between response error and confidence wedge size. T-tests on the output beta coefficients from single-subject GLMs modelling confidence width showed a significant effect of response error (*t(*19) = 7.29, *p* = 3.36 * 10^-7^). This reflects that, across subjects, trials with higher response error were associated with larger confidence wedges (lower confidence). In this analysis, there was no significant interaction of the relationship between confidence and response error by condition (*p* = 0.228).

Linear mixed-effects modelling highlighted a positive association between response error and confidence wedge size. Trials with higher response errors were associated with larger confidence wedges (*β* = 0.574, *t* = 7.741, *p* = 3.68 *10^-7^). Here, we also found an interaction of this relationship with cue type (*β* = -0.0527, *t* = -2.263, *p* = 0.0236), showing that the relationship between error and confidence was influenced by selective attention during the maintenance period. This was driven by a more positive beta (stronger association between error and confidence wedge size) in cued trials compared to neutral (see Figure 7A).

We investigated whether participants could use the continuous information in feedback to update their behaviour across trials by modelling (with a linear mixed-effects model) the relationship between the confidence error on a given trial and the change in confidence between that trial and the subsequent trial, while controlling for their confidence on the current trial (to account for regression to the mean). This showed a positive association between confidence error and changes in confidence wedge width between trials (*β* = 0.0452, *t* = 5.651, *p* = 1.64*10^-8^, shown in figure 7B). This finding indicated that participants broadened their confidence wedge (i.e., became less confident) following an over-confident trial and narrowed their confidence wedge (i.e. became more confident) following an under-confident trial.

### EEG Results

We fit a GLM (see Equation 1) to individual participants’ single-trial data, separately for each condition, with regressors to model the trial-wise response error and reported confidence.

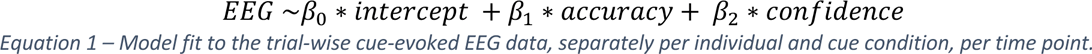

Figure 8 shows the grand-average beta weights, showing the lateralisation of alpha power evoked by the retrocue. For neutral trials, no significant clusters emerged. As expected, given the lack of spatial information in neutral cues, we saw no significant lateralisation evoked by this type of cue (Figure 8A, black line).

**Figure 8.**
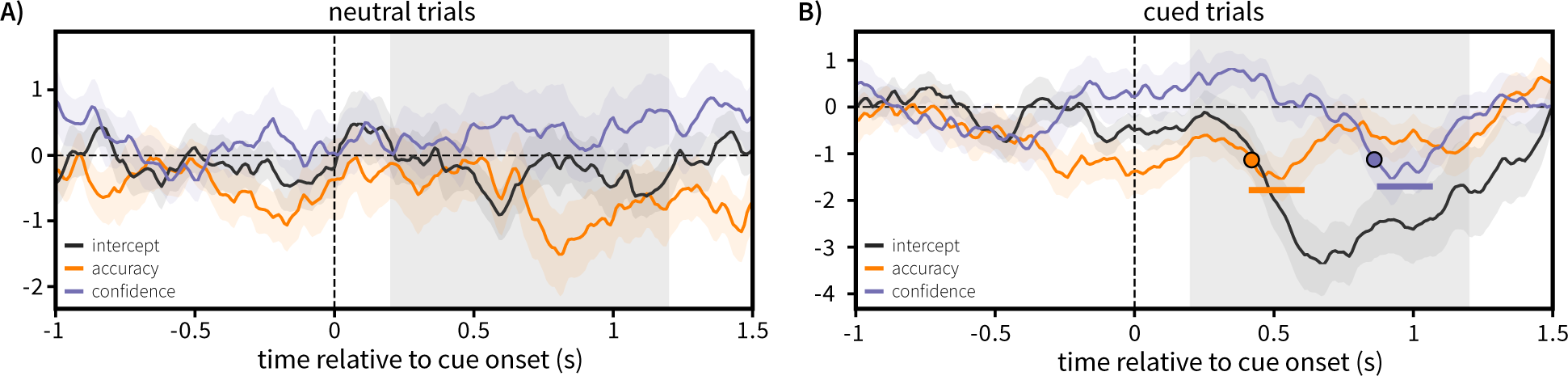
Degree of attentional selection is predictive of both objective performance and subjective uncertainty in working memory. All panels show the output of a GLM analysis of alpha lateralisation relative to the attentional cue. Models were fit separately for neutral **(A)** and cued trials **(B)**. The black line represents intercept of the model, reflecting average alpha lateralisation for the relevant cue type, controlling for both response accuracy and subjective confidence. The orange line represents the accuracy regressor, with larger values reflecting higher response accuracy (lower error). The purple line represents the confidence regressor, with larger values reflecting higher confidence (narrower reported confidence widths). Filled circles represent the jackknife estimate when time courses reached 70% of their maximum in the cluster time window. All shaded regions around solid lines represent the standard error of the mean. Solid, coloured lines above the x-axis outline time periods of the regressor time-course that survived permutation cluster testing against zero. Shaded spans of the x-axis highlight the time window used for cluster permutation testing.

We saw a significant lateralisation of alpha power after spatial retrocues (Figure 8B, black line), reflecting an orienting of attention relative to the cued item. Crucially, we observed this alpha lateralisation to be predictive of behaviour. We found that significant clusters emerged for the regressors modelling both accuracy (orange) and subjective confidence (purple). For accuracy, this negative beta reflected that increased lateralisation (relatively lower contralateral alpha power) to the cued item was associated with increased accuracy (more faithful reporting of the orientation from memory). For confidence, the negative beta reflects that increased lateralisation towards the cued item was associated with increased confidence (narrower reported confidence widths). This agrees with previous research showing that degree of internally directed attentional selection is predictive of objective performance on such working-memory tasks. Further, we extend this to show that variability in the degree of such attentional selection is also predictive of subjective working-memory confidence.

Lateralisation of alpha power appears to be related to accuracy and confidence in two qualitatively different time periods. To formally test for temporal separation in the relationship between alpha lateralisation and these aspects of behaviour, we implemented a jackknife analysis. We estimated the latencies at which the beta coefficient time courses for accuracy and confidence reached 70% of their peak value in the cluster forming window. We iteratively removed one participant from the participant pool and calculated the difference in onset latency for the two time courses, building a distribution of onset latencies, and a jackknife-based estimate of the standard error for these onset differences. This analysis revealed a significant latency difference in the peaks for the beta time courses related to accuracy and confidence (*t*(19) = -21.4, *p* = 4.70 * 10^-15^), with the confidence time course reaching threshold on average 0.443s later (mean onset latency difference 0.443 seconds, jackknife standard error 0.0927 seconds).

## Discussion

We know that individuals can introspect into the contents of their working memory, evaluating the quality of internally held information. Such introspective confidence judgements appear to track endogenous variability in working-memory performance, assumed to be related to the precision with which information is encoded into working memory. Across two experiments, we found evidence that introspective confidence judgements regarding the quality of working memories are under flexible control. Instead of being fully sealed at encoding, we provide evidence that confidence in working memory quality comes under cognitive control from multiple sources.

Across both experiments, we found behavioural evidence that orienting internal attention to selective contents in working memory improves both objective recall performance and subjective confidence estimates. Internal attention, deployed after encoding, improves both the precision of orientation recall, and the accuracy of introspective confidence judgements, with confidence more faithfully tracking the objective performance of attended memories.

In the second experiment, we show that neural markers closely associated with the orienting of internal spatial attention partially explained variability associated with both the accuracy of working-memory recall and subjective confidence. Specifically, we show that variability of the cue-induced posterior alpha lateralisation is quantitatively associated with both the accuracy of WM recall and the degree of subjective confidence in the recall of the retrieved item. Previous studies have shown that the degree of alpha lateralisation is predictive of WM accuracy across both trials (Myers et al., 2014) and across individuals (Mok et al., 2016). Here, we show that not only does the degree of attentional lateralisation towards a cued item predict objective performance but is also predictive of subjective confidence in the recalled item. Not only is subjective confidence influenced by the functional state of an item as determined by cueing, but also the degree to which neural markers of attention lateralise in response to the cue. This provides evidence that not only do subjective confidence judgements track endogenous changes in WM performance (Honig et al., 2020), but also post-encoding changes associated with attentional shifts that are indexed by ongoing cortical activity. Overall, this suggests that representations of confidence are not static, but flexibly integrate information about the functional state of the to-be-reported items.

Interestingly, we find that the degree of attentional orienting (indexed through posterior alpha lateralisation) is related to accuracy and confidence in recall in two highly temporally distinct stages. The early aspect of cue-induced lateralisation was parametrically associated with variability in objective performance, whereas the later aspect of this lateralisation was related to variability in subjective confidence. This suggests two temporally distinct processes related to how attention changes the objective and subjective aspects of WM representations, whereby early attentional selection is predictive of subsequent recall performance, and the extent of selection later on in time is related to the conscious aspect of the representation.

We also provide evidence of another important source of flexible control of subjective confidence – feedback history. Previous studies have explored within-trial relationships between working-memory performance and subjective confidence. This assumes that our subjective confidence in working-memory quality is linked solely to the individual item and unlinked to external factors beyond the trial. In our second experiment, we introduced trial-wise feedback that conveyed information about both the objective performance of WM recall and the accuracy of subjective confidence judgements. We provide evidence that subjective confidence is influenced by the accuracy of confidence judgements made on the previous trial. That is, when participants were presented with feedback that informed them they were *overconfident*, they subsequently became less confident on the next trial (and vice versa). This is at odds with hypotheses that confidence is based solely on the precision with which an item is encoded into working memory. If confidence were based solely on internal factors and agnostic to feedback, its variability across trials would be unlinked to previous trial feedback, and solely associated with current trial performance.

Taken together, these results suggest a flexible process by which we evaluate our internally held information, with introspective confidence judgements reflecting an integration of both internal and external factors. Rather than being based solely on encoding-related processes, attentional modulations of maintenance processes associated with changes in the functional state of memories can alter our introspections. Further, these introspective confidence judgements are shaped by immediate feedback history. This suggests that our belief about the working-memory contents is flexible, integrating attentional states and external feedback – creating an internal model of working-memory performance that is flexibly adjusted based on multiple information sources.

The question of what sources of information are used to construct these confidence estimates still remains. Our findings suggest an integration of information from multiple sources, but delineating their precise nature and relative contributions requires further work. Previous research has highlighted that the encoding of information into working memory is important for our introspective confidence. Our work provides convincing evidence that the degree to which items are maintained is also important, along with objective feedback regarding the accuracy of confidence estimates. Dissociating the influence of distinct memory processes is important to gaining a more holistic view of what it is that shapes our qualitative experience of memory.

In our study, we focus specifically on the maintenance period. We demonstrate that selective attention to memoranda in an internal space improves the quality of memories and confers improvements in confidence. A natural question follows: do different kinds of attentional selection affect confidence in similar ways? Other work has also shown that retrocues can also direct attention to features (or feature dimensions) of objects contained in memory (Niklaus et al., 2017) and contrasted item-based and feature-based selection directly (Hajonides et al., 2020). Integrating confidence judgements into working-memory tasks that investigate feature-based selection could help inform whether the effect of attention on confidence is driven by a spatial, item-specific reactivation, or related to the selection of the to-be-reported feature without requiring spatial information.

In this study, we deliberately did not reward participants based on the confidence judgements that they made. There is an argument that rewarding participants based on the error in their confidence judgement (as in Honig et al., 2020) generates a trade-off that encourages accuracy in their responses – in this reward strategy, participants receive reduced numbers of points where the confidence interval is much wider than required to capture the target orientation. However, the prospect of reward on each trial can have attentional effects that may influence the endogenous characteristics of working memory – for example, generating stronger trial history effects than expected following erroneous trials due to also accruing fewer points. We aimed to look at endogenous insight into working memory in the absence of trial-wise reward-based motivation to determine the effect of attention without this potential extra contribution. However, whether and how cue effects are altered in the presence or absence of reward is an interesting question. Additionally, how reward might change the metacognitive effects we observed remains an open question to investigate.

In sum, across two experiments, we show that people can make metacognitive judgements about the contents of one’s own memory, and that these judgements are related to the attentional state of the to-be-reported item. Using concurrent EEG measurements, we found distinct neural stages of processing related to attentional changes in accuracy and confidence of working memories, suggesting partially independent attentional impact. We also provide evidence that participants track confidence across trials, with beliefs about the accuracy of confidence judgements flexibly updated over time. Confidence judgements on the contents of working memory provide a rich measure of behaviour that could better enable us to investigate fluctuations in memory performance and attention, and how people combine information about memory uncertainty when utilising memories for goal-directed behaviours.

## Acknowledgements

This research was funded by a Wellcome Trust Senior Investigator Award (104571/Z/14/Z) and a James S. McDonnell Foundation Understanding Human Cognition Collaborative Award (number 220020448) to A.C.N., a BBSRC AFL Fellowship (BB/R010803/1) and the European Research Council (grant FORAGINGCORTEX, project number 101076247) to N.K., a Glyn Humphreys Scholarship to S.R.C., and supported by the NIHR Oxford Health Biomedical Research Centre. The Wellcome Centre for Integrative Neuroimaging is supported by core funding from the Wellcome Trust (203139/Z/16/Z). The funders had no role in study design, data collection and analysis, decision to publish, or preparation of the manuscript. Views and opinions expressed are those of the author only and do not necessarily reflect those of the funders. Neither the European Union nor the granting authority can be held responsible for them. We thank Jill O’Reilly, Daryl Fougnie, Sanjay Manohar and Dejan Draschkow for helpful discussions along the way. For the purpose of open access, the authors have applied a CC-BY public copyright license to any Author Accepted Manuscript version arising from this submission.

## Supplementary Analyses

### Simulation Methods

For all simulations, a standard procedure was used to generate artificial participant data. A mixture model was fitted to the participant data from experiment 1 (Bays, Catalao & Husain, 2009). This model decomposes participant response distributions as a mixture of component distributions that reflect different processes. The model fit gives rise to three parameters: *Kappa*, the concentration of the best-fitting von mises distribution that describes response variability; *Pn*, the probability of non-target responses, and; Pu, the probability of random responses (a uniform distribution that captures random guessing). This model was fit separately for cued and neutral trials per participant.

The across-subject mean and standard error *Kappa* parameter for each condition were used as parameters for a normal distribution, from which we randomly sampled to obtain a simulated precision of the response error distribution for neutral and cued trials. Using these simulated concentration parameters, we generated a response error distribution for neutral and cued trials with mean 0 and a concentration parameter dependent on the sampled value.

These simulations were performed under the null hypothesis that a perfect observer has no insight into the single-trial noise of their mnemonic representations, and thus no insight into their single-trial error - an optimal agent with perfect knowledge of the inaccuracy of their memory would adjust their response to account for this, and thus make a response with zero error on every trial. Instead, a perfect observer may have perfect insight into their across-trial error distribution, and thus their confidence estimate would be a noisy (random) sampling of this error distribution. To simulate confidence under this null distribution, we randomly sampled confidence on each trial from the same underlying distribution as response error, taking the absolute value of this sampled error (as you cannot have negative confidence).

For each simulation, an experimental dataset was simulated. We generated twenty artificial participants per simulated dataset, with 128 trials per cue condition. Ten-thousand simulated datasets were created using this procedure, allowing us to build up a simulated null distribution of the relevant test statistic under the null hypothesis that confidence estimates are generated with no insight into single-trial memory error.

### Permutation methods

An alternative method to investigate this null belief is to use permutations. Rather than simulating datasets, the experimental data are randomly shuffled (permuted) to preserve individual subject response error and confidence distributions, but remove any relationship between single-trial performance and confidence. This, across permutations, builds up a distribution of the relevant test statistic for the experimental dataset, reflecting the distribution of test statistics that arise in the data due to random chance. The proportion of test statistics, in this permuted null distribution, larger than the observed experimental effect, reflects the p-value for our observed results.

### 1 – Observed correlations between error and confidence do not arise due to chance

We show evidence that subjective confidence judgements are associated with response error, suggesting insight into performance on the individual trial. However, across trial relationships could arise not due to insight into the single-trial response error, but instead from perfect insight into the across trial response error distribution. In such a model, individuals would not have insight into their individual trial WM error as an observer with insight into the exact error in their WM representation would alter their response to account for this, leading to zero error. Instead, an individual may have perfect knowledge of the across-trial error distribution, but no insight on the individual trial. In such a scenario, confidence estimates would be generated by randomly sampling from the across-trial error distribution, to generate a subjective uncertainty in each trial. The across trial confidence distribution would then match identically the response error distribution.

To test for such an explanation for our observed results, we ran a permutation-based analysis on the experimental data gathered in experiment 1. For each subject, confidence was shuffled across trials for each cue condition separately. Spearman’s rank correlation was calculated for the relationship between response error and this shuffled confidence, separately for each cue condition. This was repeated 10,000 times for each subject, generating a null distribution of the spearman’s rank correlation between response error and confidence for each participant and condition in the task (see Supplementary Figure 1A for an example null distribution for a participant).

To test whether the observed correlation was reliably larger than the relationship expected under the null (the mean correlation of the permuted null distribution), a paired-samples t-test was performed, across subjects, on the fisher-transformed correlation coefficients, separately for each cue condition. This test was significant for both neutral trials (*t*(19) = 7.27, *p* = 3.39 x 10^-7^) and cued trials (*t*(19) 6.29, *p* = 2.43 x 10^-6^). This suggests that participants reported confidence across trials was related to response error, and integrated information about performance at the level of the individual trial. Individual participant correlation coefficients are visualised in Supplementary Figure 1B, along with summary statistics of the generated null distributions.

**Supplementary Figure 1.**
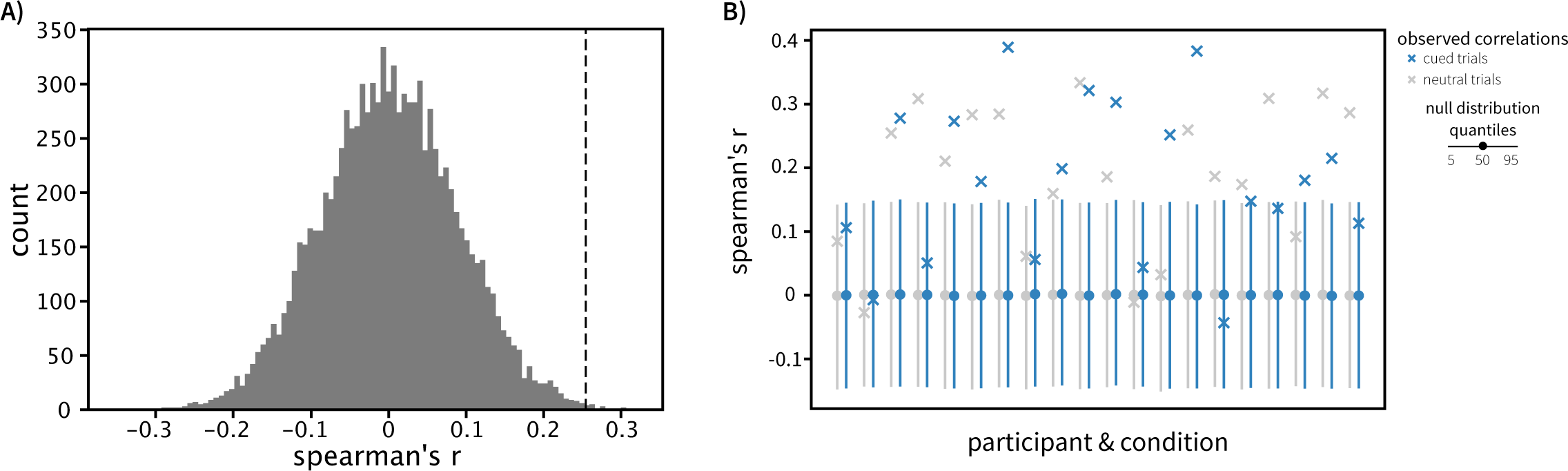
A) Histogram visualising an example permuted null distribution of correlation coefficients. Dashed black line shows the observed correlation for this participant. B) Visualisation of the observed correlations for each subject, and the relevant permuted null distribution. Solid circles show the expected (mean) permuted spearman’s rank correlation for each participant and condition. Solid lines represent the 5^th^ to 95^th^ percentile of the permuted null distributions. Crosses reflect the observed spearman’s rank correlation, separately for each condition and participant. Blue marks reflect the relevant statistics for cued trials, with grey representing neutral trials.

### 2 – Observed random effects do not arise due to perfect insight to across-trial error distributions

#### I. Permutation

To determine whether the result of our random-effects analysis (showing that the relationship between response error and confidence is modulated by cue type) was due to chance, we conducted a permutation-based control analysis. In each permutation, we shuffled the reported confidence of each participant, separately per participant and condition, to preserve the individual subjects’ confidence distributions but unlink error and confidence at the level of the individual trial. We then ran the random effects analysis, implementing the same single-subject GLM as in the main analysis, and conducting a two-sided t-test on the interaction term across participants. This t-statistic for the interaction was generated for each of the 10,000 permutations that were implemented on the data. The significance of the observed, experimental results was calculated by the proportion of permutations giving rise to a t-value larger than we observed in analysis of the experimental data (see Supplementary Figure 2A). The p-value for our observed t-value was 0.0034 (only 34 of 10,000 permutations gave rise to a larger t-value than our experimental result).

#### II. Simulation

A simulation was run to explore the range of effects that could arise under the previously stated null belief, to determine whether simulated results are concordant with the findings of our permutation-based control analysis. We simulated 10,000 experimental datasets, implementing our random-effects analysis on each experimental dataset. A distribution of the resulting test statistics (see Supplementary Figure 2B) was generated, and our experimental analysis result was compared to this distribution. We found that only 32 of 10,000 simulated datasets gave rise to a t-value larger than our experimental analysis (*p* = 0.0032).

**Supplementary Figure 2.**
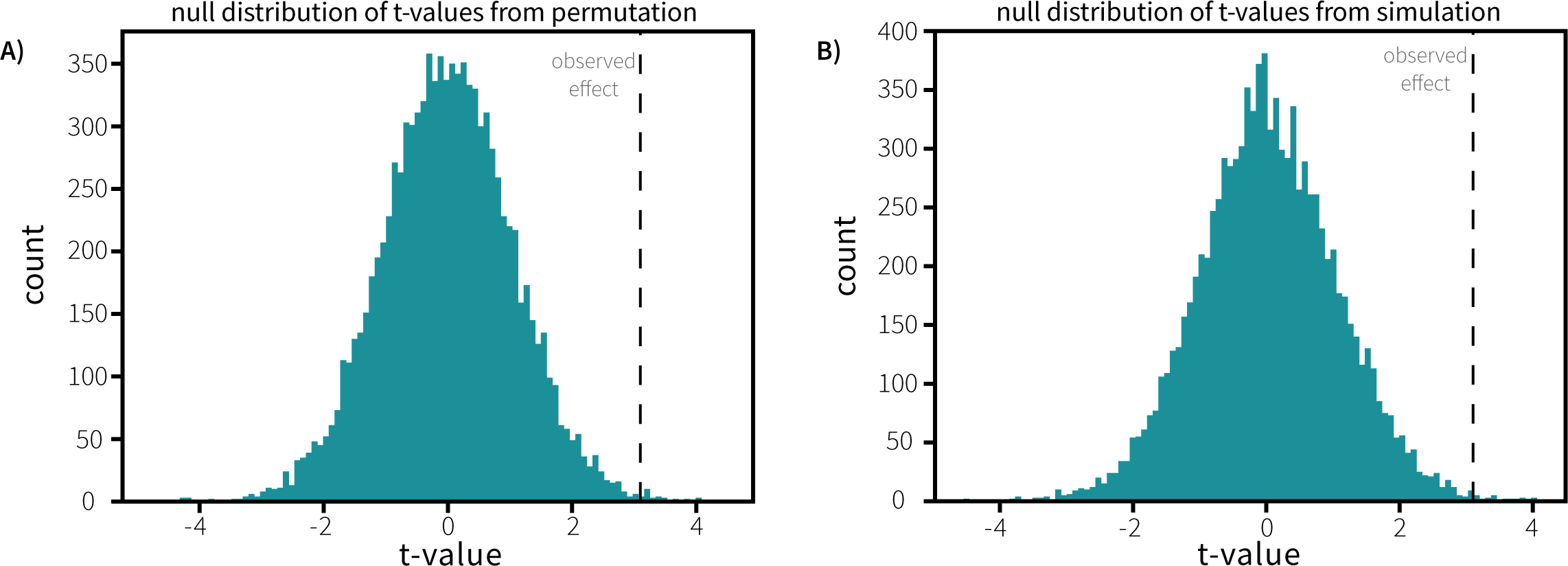
A) Distribution of t-values from permutation-based control analysis of the random effects. B) Distribution of t-values from control analysis using simulation of random effects across simulated datasets. Dashed lines in both panels represent the empirically-observed t-value for the random-effects analysis.

### 3 – Observed linear mixed-effects do not arise due to perfect insight into across-trial error distributions

#### I. Permutation

To determine whether the result of our linear mixed-effects analysis (showing that the relationship between response error and confidence is modulated by cue type) was due to chance, we conducted a permutation-based control analysis. In each permutation, we shuffled the reported confidence of each participant, separately per participant and condition, to preserve the individual subjects’ confidence distributions but unlink error and confidence at the level of the individual trial. We then ran the linear mixed-effects analysis, implementing the same model structure as in the main analysis. The t-value for the interaction term of this analysis was then generated for 10,000 permutations of the data. The significance of the observed, experimental results can be assessed by the proportion of permutations giving rise to a t-value larger than we observed in analysis of the experimental data (see Supplementary Figure 3A). The p-value for our observed t-value was 0.0014 (only 14 of 10,000 permutations gave rise to a larger t-value than our experimental result).

#### II. Simulation

A simulation was run to explore the range of effects that could arise under the previously stated null belief, to determine whether simulated results are concordant with the findings of our permutation-based control analysis. We simulated 10,000 experimental datasets, implementing our linear mixed-effects model on each experimental dataset. A distribution of the resulting test statistics (see Supplementary Figure 3B) was generated, and our experimental analysis result was compared to this distribution. We found that only 12 of 10,000 simulated datasets gave rise to a t-value larger than our experimental analysis (*p* = 0.0012).

**Supplementary Figure 3.**
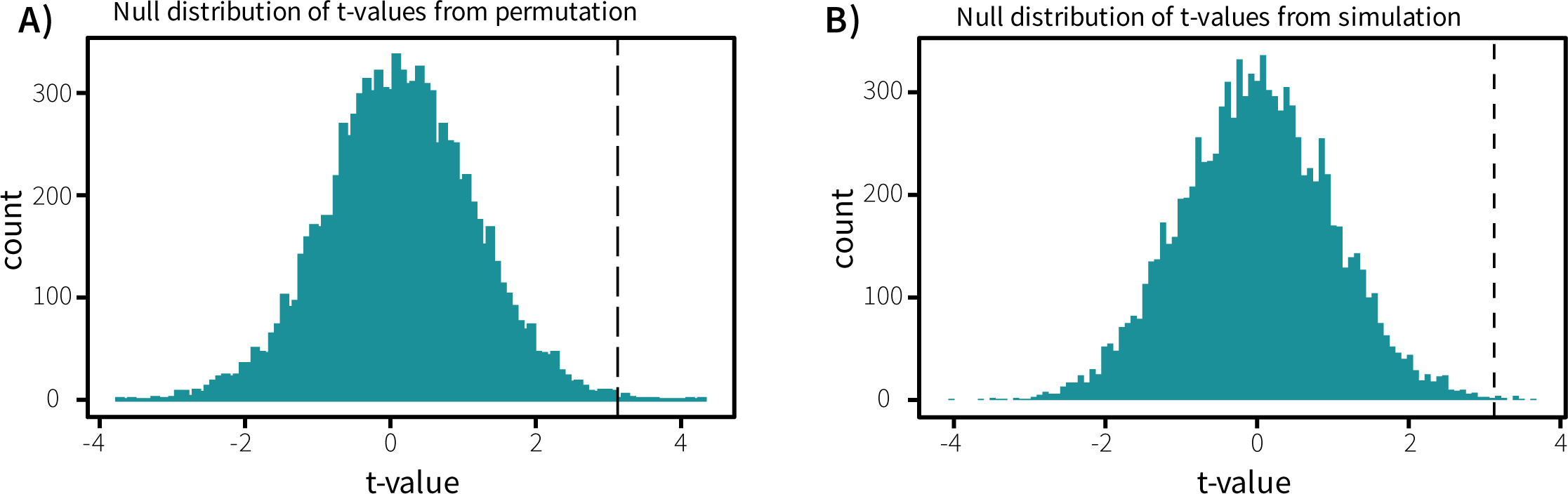
A) Distribution of t-values from permutation-based control analysis of the linear mixed-effects analysis. B) Distribution of t-values from control analysis using simulation of linear mixed-effects across simulated datasets. Dashed lines in both panels represent the empirically-observed t-value for the random-effects analysis.

